# An NMR-based biosensor to measure stereo-specific methionine sulfoxide reductase (MSR) activities *in vitro* and *in vivo*

**DOI:** 10.1101/2020.05.14.092957

**Authors:** Carolina Sánchez-López, Natalia Labadie, Verónica A. Lombardo, Franco A. Biglione, Bruno Manta, Reeba S. Jacob, Vadim N. Gladyshev, Salim Abdelilah-Seyfried, Philipp Selenko, Andres Binolfi

**Author notes:** These authors contributed equally to this work (Carolina Sánchez-López, Natalia Labadie).

## Abstract

Oxidation of protein methionines to methionine-sulfoxides (MetOx) is associated with several age-related diseases. In healthy cells, MetOx is reduced to methionine by two families of conserved methionine sulfoxide reductase enzymes, MSRA and MSRB that specifically target the *S*- or *R*-diastereoisomers of methionine-sulfoxides, respectively. To directly interrogate MSRA and MSRB functions in cellular settings, we developed an NMR-based biosensor that we call CarMetOx to simultaneously measure both enzyme activities in single reaction setups. We demonstrate the suitability of our strategy to delineate MSR functions in complex biological environments that range from native cell lysates to zebrafish embryos. Thereby, we establish differences in substrate specificities between prokaryotic and eukaryotic MSRs and introduce CarMetOx as a highly sensitive tool for studying therapeutic targets of oxidative stress-related human diseases and redox regulated signaling pathways. Our approach further extends high-resolution in-cell NMR measurements of exogenously delivered biomolecules to an entire multicellular organism.

---

Oxidation of methionine sidechains is a hallmark of cellular ageing and oxidative stress.^[1]^ Methionine oxidation produces methionine-sulfoxides (MetOx), with a chiral center at the sulfur atom giving rise to two diastereoisomers, designated *R*- and *S*-MetOx (Figure S1 a).^[2]^ Under physiological conditions, methionine sulfoxides are reduced by two families of conserved enzymes. Class A methionine sulfoxide reductases selectively act on *S*-MetOx (i.e. MSRAs), whereas class B enzymes reduce *R*-MetOx (i.e. MSRBs).^[3]^ Both MSR families are ubiquitously expressed in bacteria, yeast, plants and animals. In addition, bacteria and yeast harbor a specialized enzyme, f-*R*-MSR that only reduces free *R*-MetOx amino acids while the existence of a corresponding f-*S*-MSR has been proposed but not confirmed.^[4]^ Recently, two membrane-associated, molybdopterin-containing proteins with methionine sulfoxide reductase activity have been identified in bacteria.^[5]^

MSRs are implicated in the development of age-related neurodegenerative^[6]^ and cardiovascular^[7]^ disorders. Evolutionary conservation of MSRs and their selective substrate specificities further suggest that methionine oxidation and reduction may convey cellular signaling activities, similar to reversible phosphorylation of serine, threonine and tyrosine residues.^[8]^ Regulatory roles of methionine oxidation have recently been discovered in bacteria^[9]^ and yeast.^[10]^ In mammals, reversible Met-oxidation appears to regulate actin polymerization^[11]^ and Ca^2+^/Calmodulin-dependent kinase II (CamKII) activity.^[12]^ These observations stimulated renewed interest in MSRs and prompted the development of analytical tools to specifically measure MSRA or MSRB activities *in vitro*^[13]^ and *in vivo*, including by fluorescently labeled Met sulfoxide derivatives^[14]^ or MSRA/B-YFP chimeras.^[15]^ While these are useful tools for single reductase assays, they do not allow to discriminate between MSRA and MSRB activities in the same reaction. In addition, they are unsuitable for the characterization of f(*R*/*S*)MSR activities because of the single amino-acid requirements of these enzymes, which preclude substrate derivatization.^[4a]^ To overcome these limitations, we developed an NMR-based biosensor to monitor native MSR activities in different biological environments. Our approach is based on the unique sensitivity of the NMR chemical shift and its ability to stereo-specifically report on methionine oxidation states in complex mixtures such as reconstituted reductase reactions and cell extracts with endogenous MSR activities.^[16]^ Furthermore, microinjection of a newly synthesized biosensor, that we called CarMetOx, into zebrafish embryos and direct NMR acquisition on those samples allowed us to characterize MSRA and MSRB activities simultaneously *in vivo*. This expands the repertoire of existing in-cell NMR methodologies for the detection of exogenously delivered, isotopically enriched biomolecules in cultured cells^[17]^ to a complex vertebrate organism.

To define the experimental basis for our approach, we initially chose the intrinsically disordered protein gamma-synuclein (γ-Syn) and free L-methionine as model MSR substrates. First, we oxidized, recombinant γ-Syn and L-Met to their corresponding sulfoxides.^[16]^ 2D ^1^H-^15^N SOFAST-HMQC NMR spectra of reduced and oxidized ^15^N isotope-enriched γ-Syn revealed discrete chemical shift changes for Met1 and Met38, indicating that no major structural changes occurred and that no other residues were modified (**Figure S1 b-d**). We detected two well-resolved NMR resonances for Tyr39, which likely reported on the *S*- and *R*-diastereoisomers of neighboring Met38Ox. Similarly, 2D ^1^H-^13^C HSQC spectra of ^15^N/^13^C-labeled, oxidized L-methionine (MetOx) displayed pronounced splitting of the ^1^Hγ-^13^Cγ cross-peak (**Figure S1 e**). Time-resolved NMR analysis of γ-Syn and Met oxidation with H_2_O_2_ revealed the formation of equimolar *S*- and *R*-diastereoisomers, with similar second-order rate kinetics (**Figure S1 f**).^[18]^ Next, we expressed and purified recombinant yeast MSRA and MSRB and reconstituted individual reductase reactions with isotope-enriched, oxidized γ-Syn and L-MetOx. To delineate enzyme activities, we followed γ-Syn Tyr39 and MetOx Hγ-Cγ signals over time.^[19]^ Both enzymes produced characteristic changes in the NMR spectra of γ-SynOx and MetOx that clearly reflected the stereospecific reductions of the respective substrates (**Figure 1 a**). Reconstituted MSRA-MSRB experiments further allowed us to unambiguously assign cross-peaks corresponding to *S*- or *R*-MetOx isomers of γ-SynOx and MetOx. Time-course analysis revealed that MSRA activity on *S*-γ-SynOx was slightly higher than for *S*-MetOx, whereas MSRB reduced *R*-γ-SynOx but not free *R*-MetOx (**Figure 1 b** and **Table S1**), in agreement with previous data about the substrate specificities of yeast MSRA and MSRB.^[20]^

**Figure 1.**
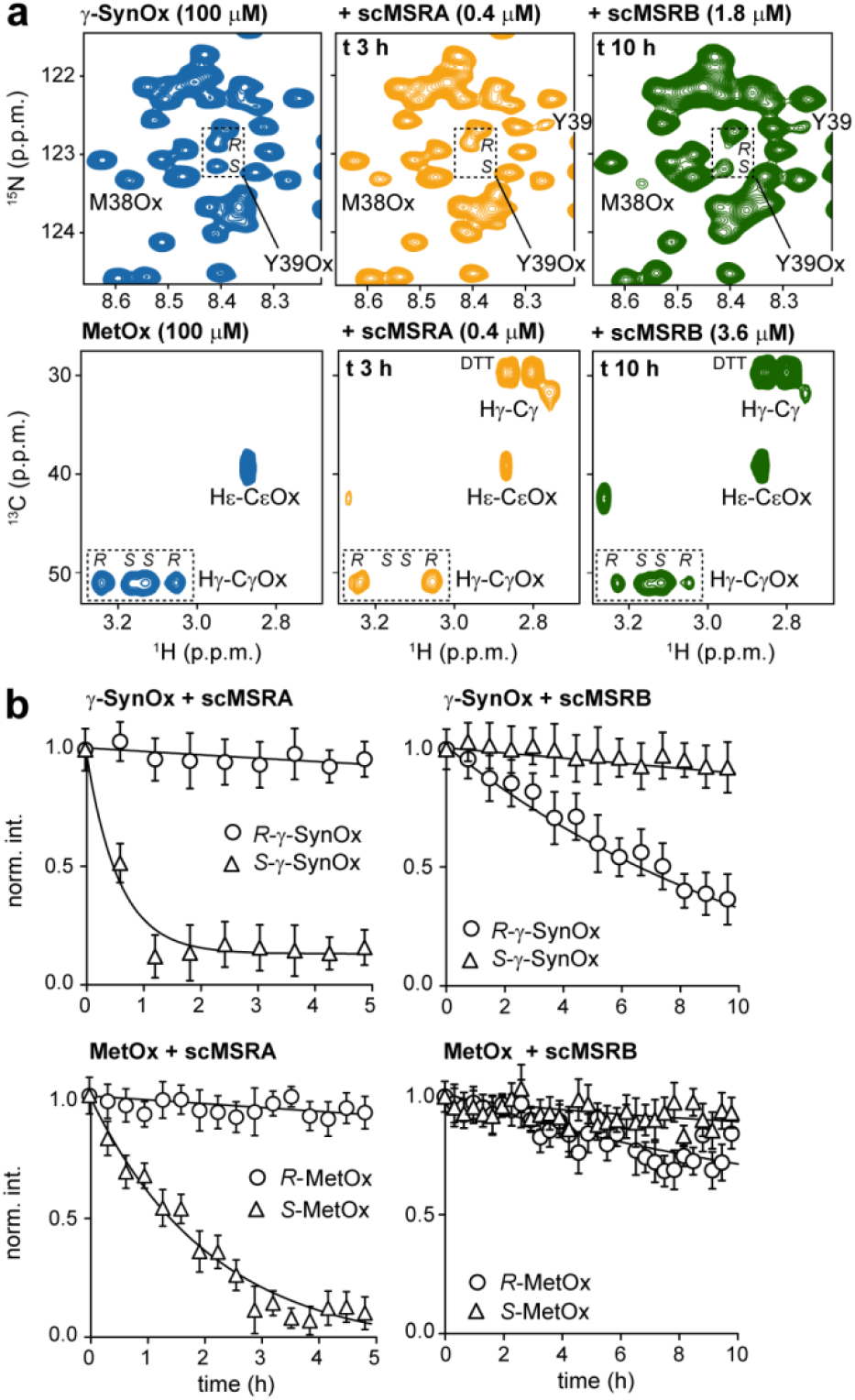
Time-resolved MSRA and MSRB activities. (a) ^1^H-^15^N SOFAST-HMQC spectra of ^15^N isotope-enriched γ-SynOx (top) and ^1^H-^13^C HSQC spectra of ^13^C isotope-enriched MetOx (bottom) without (blue) and with MSRA (yellow) or MSRB (green) at the indicated incubation times. Dotted boxes highlight cross-peaks that report on *S*- and *R*- stereoisomers of neighbouring methionine-sulfoxides. (b) Time-resolved NMR profiles of sulfoxide reduction of γ-SynOx (top) and MetOx (bottom) by MSRA (left) or MSRB (right).

While γ-SynOx and MetOx model substrates may be used to characterize MSRA and MSRB activities *in vitro*, their suitability as general MSR reporters in live cells or *in vivo* is limited. Free amino acids such as L-Met are rapidly incorporated into proteins or co-factors, including S-adenosyl-Met. Similarly, NMR detection of *R*- and *S-* signals of intracellular ^15^N-γ-SynOx is hampered by fast relaxation and physiological protein turnover.^[21]^ To overcome these drawbacks, we engineered a synthetic MSR reporter based on L-Met as a scaffold, for which the detection of *R*- and *S*-diastereoisomers is straightforward. To this end, we derivatized ^15^N-^13^C isotopically enriched L-Met with diethyl-pyrocarbonate (DEPC), a reagent that modifies the α-amino group of Met^[22]^ into a carbethoxylated L-Met compound, which we refer to as ‘CarMet’ (**Figure 2 a** and **Figure S2**). The CarMet reaction produces an L-Met amide proton that can be detected in 2D ^1^H-^15^N SOFAST-HMQC^[23]^ experiments with good signal-to-noise (**Figure 2 b**). Oxidation of CarMet to CarMetOx produces two well-resolved cross-peaks of the racemic mixture of *S*- and *R*-diastereoisomers. By reacting CarMetOx with either MSRA or MSRB, we assigned *S*- and *R*-resonances (**Figure 2c**). Furthermore, time-course experiments showed that CarMetOx fully recapitulated the behavior of an oxidized protein substrate rather than a free amino-acid (**Figure 2d** and **Table S1**).

**Figure 2.**
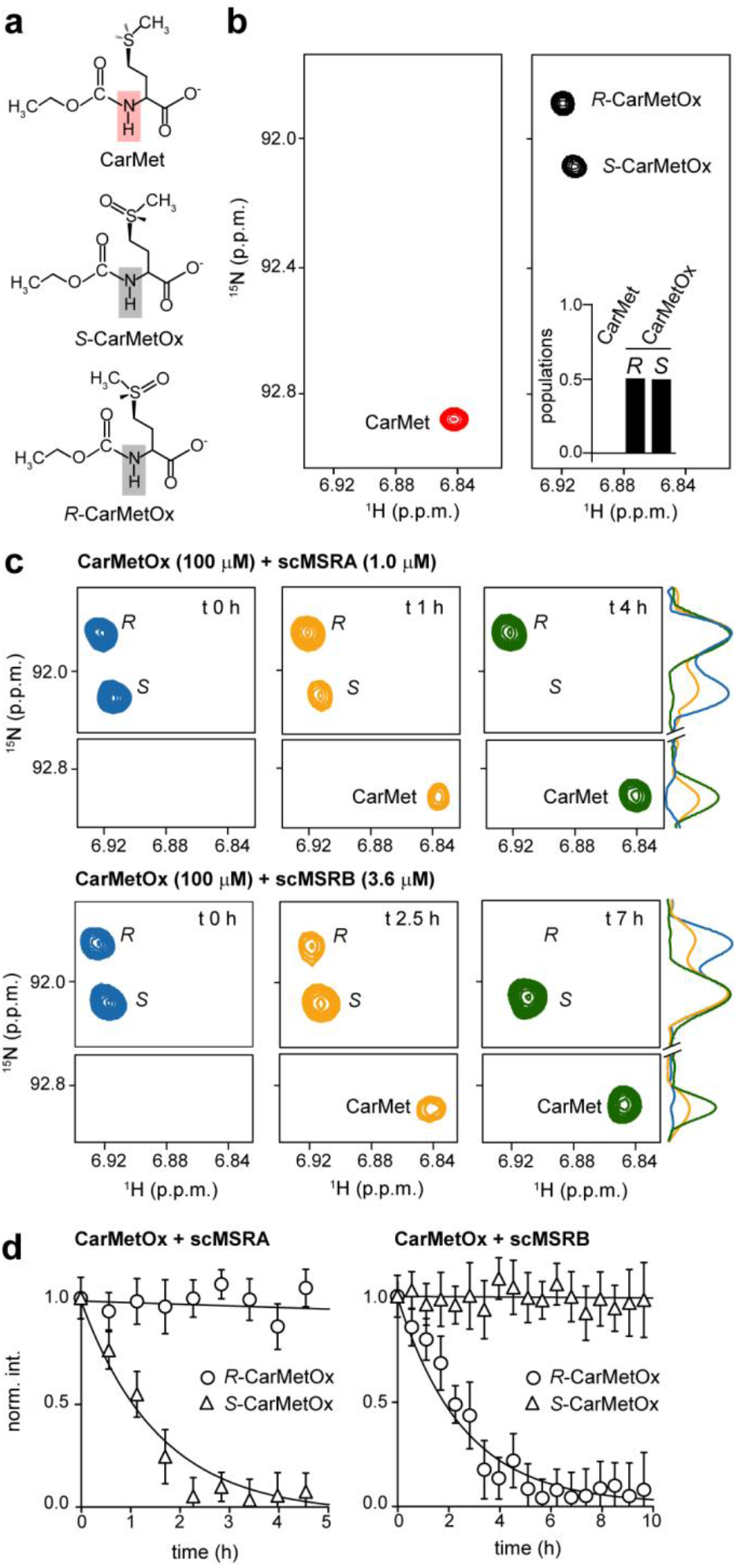
CarMetOx as a biosensor for MSRA and MSRB activities. (a) Chemical structures of CarMet and *R*- and *S*-CarMetOx. The amide groups of each compound observed in ^1^H-^15^N correlation spectra are highlighted. (b) ^1^H-^15^N SOFAST-HMQC spectra of 500 μM CarMet (red) and CarMetOx (black). Two sets of NMR signals in the CarMetOx spectrum correspond to the racemic mixture of *S*- and *R*-diastereoisomers. Inset shows the NMR quantification of each species. (c) NMR monitoring of CarMetOx reduction by MSRA (top) or MSRB (bottom). 1D NMR spectra on the right correspond to the ^15^N traces of the 2D spectra at 6.92 p.p.m. in the ^1^H frequency. (d) Time-resolved NMR profiles of CarMetOx reduction by MSRA (left) and MSRB (right).

Next, we asked whether we can use CarMetOx to quantify native MSRA and MSRB activities in complex physiological environments of cell lysates. We added ^15^N isotope-enriched CarMetOx to bacterial and mammalian lysates and recorded consecutive ^1^H-^15^N SOFAST-HMQC NMR experiments (**Figure 3a** and **Figure S3a**). We measured progressive reductions of *S*- and *R*-CarMetOx resonances in 2D NMR spectra and the corresponding build-up of single CarMet signals, which enabled us to directly monitor cellular MSRA and MSRB activities in both mixtures.

**Figure 3.**
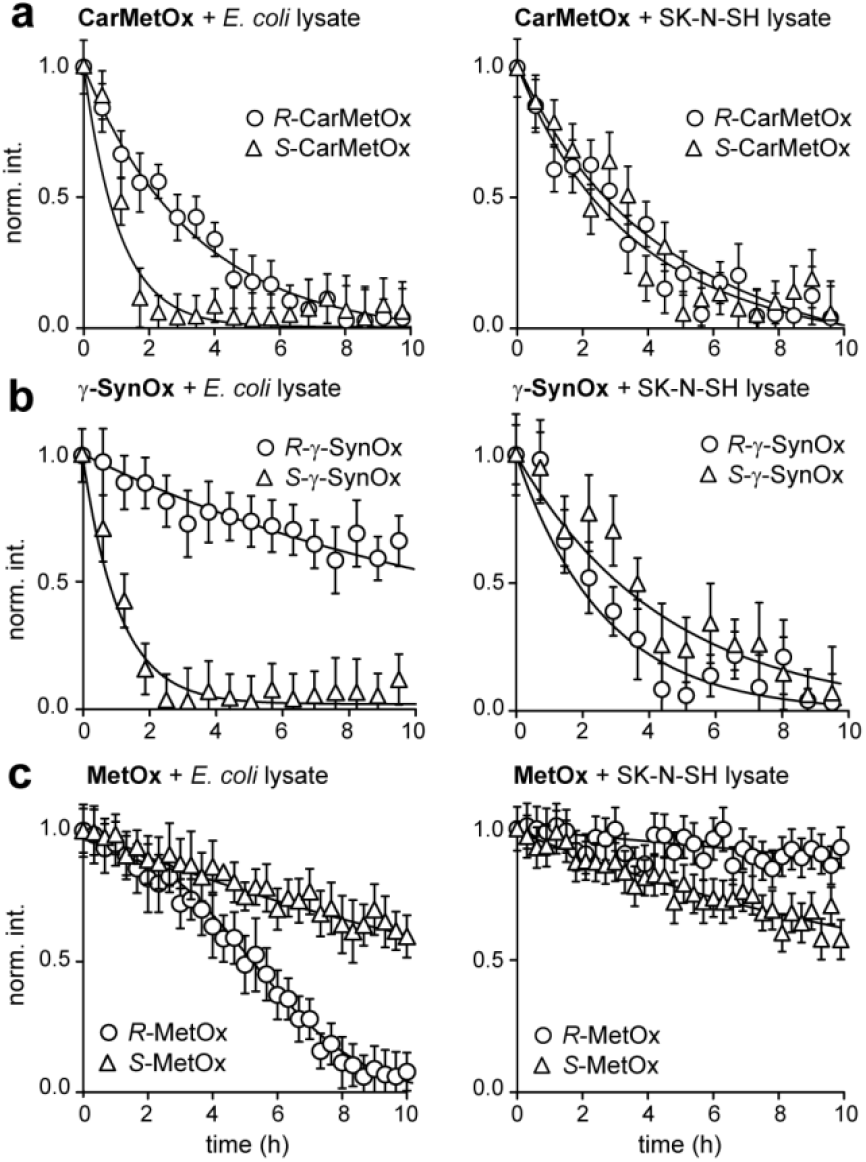
Stereo-specific reduction of Met sulfoxides in bacterial and mammalian cell lysates. Time-resolved NMR profiles of sulfoxide reduction by endogenous MSRs on isotope-enriched CarMetOx (a), γ-SynOx (b) and MetOx (c) in *E. coli* (left) and SK-N-SH (right) cell lysates. Experiments were performed with 100 μM substrates concentrations and 4.0 mg/mL of total lysate protein.

In a second step, we repeated these experiments with free MetOx and γ-SynOx to assess amino-acid and protein-specific differences of MSR activities in these environments. We found that reduction of γ-SynOx *S*-diastereoisomers proceeded slightly faster in *E. coli* versus mammalian SK-N-SH cell lysates, whereas rates of *S*-MetOx reduction were similar in both types of lysates (**Figure 3b, c**). These results suggested that bacterial and mammalian MSRAs exhibit comparable overall activities and substrate specificities. Moreover, cellular MSRAs processed γ-SynOx *S*-sulfoxides more efficiently than in MetOx, in agreement with previously reported MSRA preferences for disordered protein substrates.^[24]^ Mammalian MSRBs displayed higher activities towards γ-SynOx *R*-sulfoxides compared to prokaryotic enzymes (**Figure 3b**), which may reflect the greater abundance and diversity of MSRBs in higher organisms.^[25]^ Interestingly, *R*-diastereoisomers of free MetOx were efficiently reduced in *E. coli* but not in mammalian cell lysates (**Figure 3c**). This observation suggested that other enzymes, probably f-*R*-MSRs, targeted *R*-MetOx in bacterial lysates and that such activities were not present in lysates of mammalian SK-N-SH cells.^[25]^ Furthermore, we measured comparable rates of *S*-MetOx reduction in *E. coli* and SK-N-SH lysates, arguing against additional f-*S*-MSR activities in this prokaryote (**Figure 3c**) and that MSRA-type enzymes may be sufficient to maintain methionines in their reduced states in bacteria. Together, these results established that time-resolved NMR measurements of MSR substrates with different characteristics provided mechanistic insights into endogenous MSRA and MSRB activities in bacterial and human cell lysates allowing also the deconvolution of individual contributions within the large MSR family. Furthermore, we showed that CarMetOx is an excellent substrate to characterize MSR activities in complex environments as its enzymatic reduction is easily detected in both prokaryotic and eukaryotic cell lysates.

Finally, we set out to explore the possibility of using CarMetOx for studying MSRA and MSRB activities in a living organism. Inspired by earlier *in vivo* NMR metabolmics studies in live multicellular organisms,^[26]^ and in-cell NMR experiments in *Xenopus laevis* oocytes,^[27]^ we microinjected isotope-enriched CarMetOx into developing zebrafish embryos (*Danio rerio*). Delivery of water soluble compounds into the yolk of one-cell stage embryos results in efficient targeting to blastoderm cells by cytoplasmic streaming, a well-established phenomenon in developing fish (**Fig. 4a**).^[28]^ We confirmed blastoderm localization of yolk-injected CarMetOx by fluorescence microscopy of a fluorescein isothiocyanate (FITC)-tagged L-Met analogue (**Fig. 4b** and **Fig. S4a**). Next, we injected ^15^N isotope-enriched CarMetOx into zebrafish embryos, which we collected in a 5 mm Shigemi NMR tube (**Fig. 4c**). 2D ^1^H-^15^N SOFAST-HMQC spectra recorded at different time-points post injection initially revealed *R-* and *S-* NMR signals (after 0.75 h, **Fig. 4d**). Over time, *R-* and *S-* resonance intensities of CarMetOx progressively diminished, whereas the single NMR cross-peak of the reduced biosensorincreased (**Figures 4d, e**). Reduction of the CarMetOx *S*-diastereoisomers was faster than of *R*-species, suggesting that global zebrafish MSRA activities may be higher than those of MSRBs at early stages of embryonic development. We did not detect leakage cross-peaks, which confirmed that observed CarMetOx and CarMet signals originated from intact embryos (**Figure S4b**). Following NMR measurements, we transferred embryos back to E3 media and followed their development. In comparison to control animals, embryos exhibited a developmental delay while inside the NMR tube. However, normal maturation resumed after NMR experiments, when embryos were put into fresh medium. More than 65 % of NMR specimens matured into healthy and morphologically indistinguishable fish, which established that CarMetOx injections and NMR measurements did not negatively impact their overall development (**Figures 4 f, g**). To gain further insights about cellular MSRA and MSRB activities, we repeated CarMetOx NMR experiments in lysates that we prepared from 512-cell and 26-somite stage embryos (**Figure S5**). Whereas we clearly detected CarMetOx to CarMet reduction in these mixtures, overall repair rates were greatly reduced. Notably, we had previously observed similarly compromised turnover effects with unrelated oxidized protein substrates in mammalian cell lysates.^[16]^ The most plausible explanation for these differences between in-cell and in-lysate NMR experiments relate to the much higher dilution of cytoplasmic factors in lysates compared to intact cells. Accordingly, effective MSR concentrations are smaller in lysates than in intact embryos.

**Figure 4.**
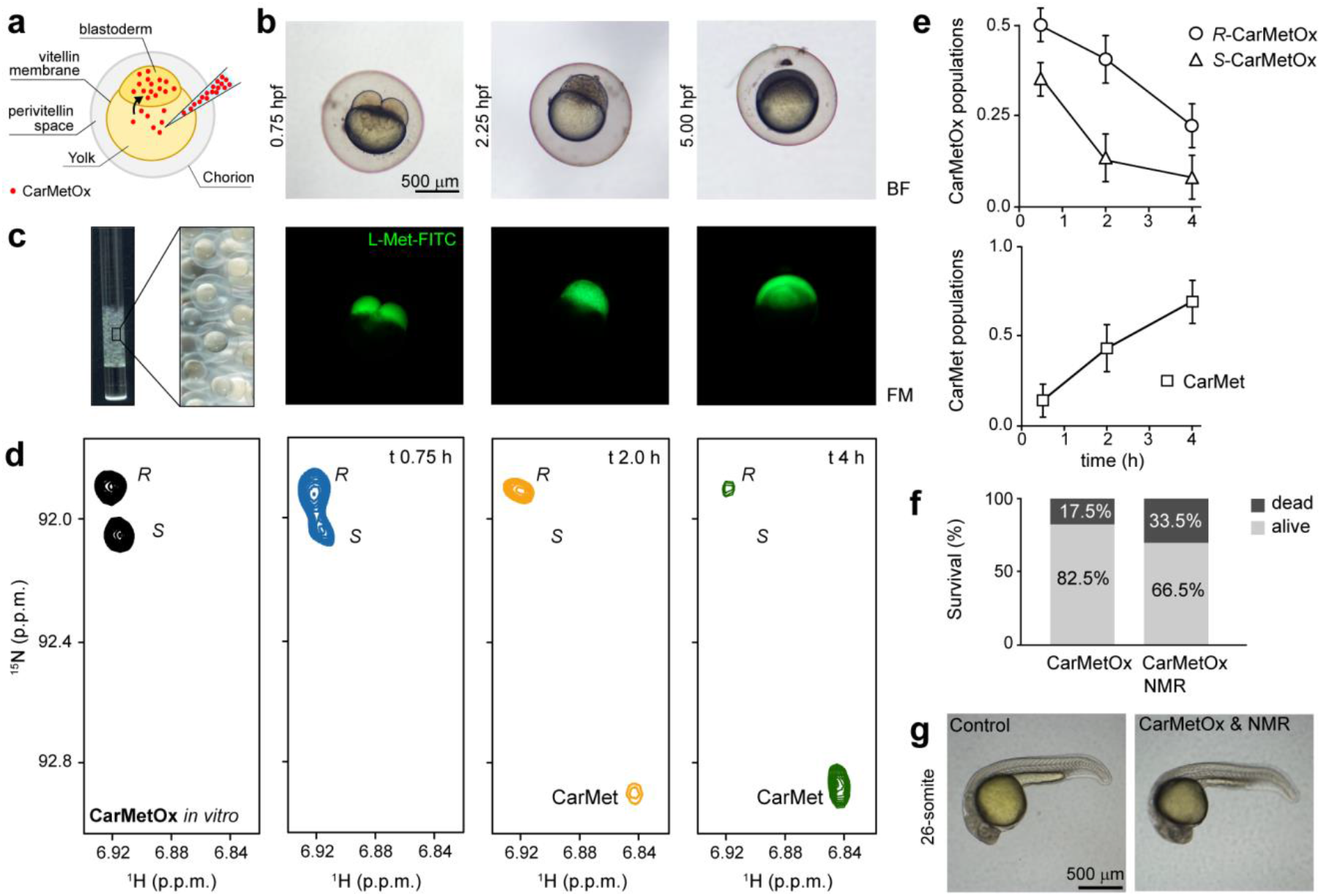
CarMetOx as a biosensor for MSRA and MSRB activities in zebrafish embryos. (a) Schematic depiction of CarMetOx microinjection in the yolk of one-cell stage zebrafish embryos. (b) Bright field (top) and fluorescence (bottom) microscopy of zebrafish embryos microinjected with FITC-labelled L-Met. The developmental stage is indicated on the left. (c) Image of embryos loaded into a 5 mm NMR tube. (d) 2D ^1^H-^15^N SOFAST HMQC NMR spectra of isotope-enriched CarMetOx *in vitro* (black) and inside zebrafish embryos at the indicated hours post microinjection (blue, yellow and green). Each time-point was acquired on an independent sample (e) Quantification of relative amounts of S- and R-CarMetOx (top) and CarMet (bottom) at the different experimental time points. (f) Survival percentage of zebrafish embryos microinjected with CarMetOx (left) and microinjected with CarMetOx and subjected to NMR measurements (right) at the 26-somite stage (~24 h hours post injection). See supporting information for detailed numbers of embryo viability (g) Bright field microscopy of zebrafish embryos non-treated (left) and after CarMetOx microinjection and NMR measurements.

In summary, we introduced an NMR-based biosensor for monitoring MSR activities *in vitro*, in cell lysates and in developing zebrafish embryos. CarMetOx synthesis is straightforward and the compound is easily purified via aqueous-phase reactions. It is stable in biological samples and is not cytotoxic. In combination with time-resolved NMR measurements, the CarMetOx biosensor allows to quantify MSR activities under experimental conditions that approximate cellular *in vivo* settings, including zebrafish embryos. We believe that our results provide the experimental benchmarks for future *in vivo* NMR routines to perform structural and functional studies of microinjected biomolecules, including peptides and proteins, in a live vertebrate.

While CarMetOx is less sensitive than other existing, fluorescence-based MSR reporters^[14–15]^ it comprises the first bio-analytical tool to monitor MSRA and MSRB activities simultaneously. This may facilitate the screening of stereo-specific inhibitors of bacterial MSRs which are currently considered virulence factors in microbial infections.^[29]^ Furthermore, all methionine oxidation states of our synthetic reporter have well defined and easy to identify NMR signatures, including the reduced state (CarMet), the *R-* and *S-* sulfoxides (CarMetOx) and the sulfone (CarMetona) (**Figure 3a, b** and **Supp. Fig. 6**), suggesting that CarMet could also be used as a probe to monitor cellular oxidative stress conditions promoting protein methionine oxidation *in vivo.* Given the prominence of oxidative stress in various human diseases and the importance of MSR enzymes in oxidative damage repair, we are particularly intrigued about the prospects of using CarMetOx in assays to study the regulation of MSRA and MSRB proteins and to evaluate strategies that stimulate their activities. Such information will aid the development of new disease biomarkers and the identification of novel cellular targets for therapeutic intervention.

## Supporting information

Experimental Details and Supporting Figures and Tables

## Experimental Section

Experimental Details are available in the supporting Information associated to this work.

## Acknowledgements

A.B. and V.A.L. acknowledge CONICET, ANPCyT (PICT 2017-1241) and Fundación IBR for financial support. P.S. is a recipient of an ERC Consolidator Grant #647474 (NeuroInCellNMR). F.A.B. acknowledges CONICET for doctoral fellowship. C.S.L. acknowledges SECTEI CDMX, Mexico for a post-doctoral fellowship. We thank Dr. Luciano Abriata and Dr. François-Xavier Theillet for helpful discussions, Sebastian Graziati for assistance with fish handling and Alejandro Gago and Andrea Coscia for maintenance of the NMR facility.

