## Supplementary material for "An NMR-based biosensor to measure stereo-specific methionine sulfoxide reductase (MSR) activities *in vitro* and *in vivo*": Experimental Details and Supporting Figures and Tables

Reagents. $^{15}N$-$^{13}C$-enriched L-methionine was purchased from Cortecnec (Cortecnec, France, CCN2000P025). 30 % hydrogen peroxide ($H_2O_2$) and anhydrous glycerol were purchased from Ciccarelli Laboratories (Argentina). Diethyl pyrocarbonate (DEPC), sodium bicarbonate ($NaHCO_3$), dibasic sodium phosphate ($Na_2HPO_4$), sodium chloride (NaCl), imidazole, 1,4-dithiothreitol (DTT), deuterium oxide ($D_2O$, 99.9 %) and phenylmethylsulfonyl fluoride (PMSF) were from Sigma. Complete protease inhibitor cocktail was from Roche. Kanamycin sulfate and Streptomycin sulfate were from Applichem (Applichem ITW Reagents, Germany). Isopropyl β-D-1-thiogalactopyranoside (IPTG) was obtained from Roth (Carl Roth, Germany). All chemicals were reagent grade and used without further treatment.

Protein expression and purification. $^{15}N$ isotopically enriched gamma-synuclein (γ-Syn) was obtained by over-expressing the pT-T7 plasmid containing the DNA sequence of human γ-Syn in M9 minimal medium supplemented with 1 g/L of $^{15}NH_4Cl$ and 4 g/L D-glucose. Purification was performed using a protocol previously reported for the purification of α-syn.[1] Protein purity was assessed by SDS-PAGE and quantified spectrophotometrically using an extinction coefficient of 1490 $M^{-1}$ $cm^{-1}$ at 280 nm. Methionine-oxidized γ-syn (γ-SynOx) was obtained by incubating 750 μM of protein dissolved in MQ water with $H_2O_2$ (2 %) at $4^o$ C for 2 h.[2] The resulting solution was frozen at -80° C, lyophilized twice and resuspended in NMR buffer (20 mM sodium phosphate, 150 mM NaCl, pH 7.0, 10 % $D_2O$) to remove the excess of $H_2O_2$. Quantitative methionine oxidation was confirmed by NMR spectroscopy as previously reported.[2]

*Saccharomyces cerevisiae* MSRA and MSRB (scMSRA and scMSRB) were obtained by plasmid over-expression, pET28a_ScMSRA and pET28_ScMSRB, respectively; in *Escherichia coli* BL21 (DE3) following previously described procedures.[3] Briefly, bacteria were grown in Luria-Bertani medium (Thermo Fisher) until an $OD_{600}$ of 0.6 was reached. Overexpression was performed in the presence of 1 mM IPTG for 4 hours at 37° C (scMSRA) or overnight at 25° C (scMSRB). Cells were resuspended in lysis buffer (25 mM Tris, 1 mM EDTA, 100 mM NaCl, 10 mM imidazole, 1 mM DTT, 1 mM PMSF, pH 7.7) and sonicated in ice 5 times for 10 seconds at 30 W sonicator power (Branson, U.S.A.) with 2 minutes intervals between cycles. Lysates were cleared by centrifugation (15 min at 18000 rpm) and scMSRA and scMSRB were purified from the supernatant by affinity chromatography using Ni-NTA agarose beads, as previously outlined.[4] Purity was confirmed by SDS-PAGE. Proteins were quantified spectrophotometrically using extinction coefficients of 34630 $M^{-1}$ $cm^{-1}$ (scMSRA) and 23950 $M^{-1}$ $cm^{-1}$ (scMSRB) at 280 nm. Enzymes were used without removing the His-tag. The pure proteins were aliquoted and stored with 10 % glycerol at -80° C until use. Repeated thawing and freezing of lysate aliquots was avoided to preserve enzyme activities.

MetOx preparation. A 500 μL, 15 mM $^{15}N$-$^{13}C$-isotopically enriched L-methionine (Met) solution was prepared in MQ water and treated with 100-fold excess of $H_2O_2$. The mixture was incubated at $4^o$ C overnight, frozen at -80° C and lyophilized twice to remove non-reacted $H_2O_2$. Quantitative Met oxidation and the absence of other oxidation products, such as Met sulfone, was confirmed using NMR spectroscopy. $^1H$ and $^{13}C$ NMR chemical shifts of MetOx were in full agreement with previously published results.[5]

CarMet and CarMetOx synthesis. CarMet and CarMetOx were synthesized in aqueous solutions by incubating non-isotopically enriched or $^{15}N$-$^{13}C$ isotopically enriched L-Met with diethyl-pyrocarbonate, a reagent that efficiently adds a carbethoxy moiety to the ε- and α-amino groups of amino acids, peptides and proteins in aqueous solutions.[6] A 100 μL, 5 mM $^{15}N$-$^{13}C$-isotopically enriched L-methionine solution was prepared in fresh $NaHCO_3$ buffer (15 mM, pH 9.0) and treated with 10 equivalents of DEPC. DEPC stock solutions were prepared fresh in absolute ethanol from the pure liquid. Reaction was carried out at 4$^o$ C overnight. At this point, $^{15}N$-$^{13}C$-isotopically enriched L-methionine is quantitatively converted to CarMet (N-carbethoxyl-L-methionine) and the excess of DEPC is decomposed to ethanol and $CO_2$.[6] The reaction was then diluted to 10 mL with MQ water and treated with 100 equivalents of $H_2O_2$ at 4$^o$ C for 2h. This large dilution before oxidation is recommended to lower the concentration of $H_2O_2$ and to avoid the oxidation of CarMet to the corresponding sulfone (CarMetona) during the lyophilization process (**Figure S6**). After oxidation, the sample was frozen at -80° C and lyophilized to remove unreacted $H_2O_2$ and other reaction byproducts (ethanol, $CO_2$). The procedure results in the formation of a white powder containing a racemic mixture of Carmet *R*- and *S*-sulfoxides (CarMetOx). Under these conditions, the reaction is quantitative and does not result in the formation of other oxidation products such as CarMetona. CarMetOx was finally dissolved in MQ water at concentrations ranging from 5-25 mM and stored at -80$^o$ C until further use.

Structural characterization of L-Met, CarMet and CarMetOx along with its purity was performed by 1D $^1$H, 2D $^1$H-$^{13}$C and 2D $^1$H-$^{15}$N NMR spectroscopy (**Figure 2** and **S2**). NMR parameters were as follows: (i) L-Met: $^1$H NMR (700.20 MHz, $H_2O$/10% $D_2O$, non-isotopically enriched) δ 2.04 (s, 3H, Hε), 2.08 (m, 2H, Hβ), 2.55 (t, J = 7.91 Hz, 2H, Hγ), 3.78 (t, J = 5.99 Hz, 1H, Hα), 7.81 (s, 2H, NH); $^{13}$C NMR (176.06 MHz, $H_2O$/10% $D_2O$, $^{15}$N-$^{13}$C-isotopically enriched, from $^1$H-$^{13}$C HSQC) δ 16.78 Cε, 31.8 Cγ, 32.3 Cβ, 56.86 Cα; (ii) CarMet: $^1$H NMR (700.20 MHz, $H_2O$/10% $D_2O$, non-isotopically enriched): δ 1.15 (t, J = 7.38 Hz, 3H, Ha), 1.89 (m, 2H, Hβ), 2.03 (s, 3H, Hε), 2.50 (m, 2H, Hγ), 3.52 (m, 2H, Hb), 4.01 (m, 1H, Hα), 6.83 (d, J = 8.12 Hz, 1H, HN); $^{13}$C NMR (176.06 MHz, $H_2O$/10% $D_2O$, $^{15}$N/$^{13}$C-isotopically enriched, from $^1$H-$^{13}$C HSQC) δ 17.0 Cε, 32.64 Cγ, 34.09 Cβ, 58.47 Cα; $^{15}$N NMR (70.95 MHz, $H_2O$/10 % $D_2O$, $^{15}$N-$^{13}$C-isotopically enriched, from $^1$H-$^{15}$N SOFAST-HMQC) δ 92.8 N; (iii) CarMetOx: $^1$H NMR (700.20 MHz, $H_2O$/10 % $D_2O$, non-isotopically enriched ) δ 1.16 (t, J = 7.36 Hz, 3H, Ha), 2.07 (m, 2H, Hβ), 2.64 (s, 3H, Hε), 2.86 (m, 2H, Hγ), 3.52 (m, 2H, Hb), 4.01 (m, 1H, Hα), 6.91 (m, J = 7.88 Hz, 3.18Hz, 1H, NH); $^{13}$C NMR (176.06 MHz, $H_2O$/10 % $D_2O$, $^{15}$N/$^{13}$C-isotopically enriched, from $^1$H-$^{13}$C HSQC) δ 39.5 Cε, 52.37 Cγ, 28.0 Cβ, 58.26 Cα; $^{15}$N NMR (70.95 MHz, $H_2O$/10 % $D_2O$, $^{15}$N-$^{13}$C-isotopically enriched, from $^1$H-$^{15}$N SOFAST-HMQC) δ 91.9 N (*R*-diastereoisomer) 92.1 N (*S*-diastereoisomer).

<u>*Derivatization of L-Met with FITC.*</u> Initially, a 25 mM fluorescein isothiocyanate (FITC, Sigma F7250) stock solution was prepared in pure DMSO. 100 μL of 5 mM $^{15}$N/$^{13}$C isotopically enriched L-Met was dissolved in 50 mM $NaHCO_3$ at pH 8.5 and reacted with 2.5 mM of FITC at 4° C in the dark overnight according to the manufacturer´s protocol. We used substoichiometric amounts of FITC to minimize the amount of free FITC after the reaction. The reaction endpoint was confirmed by $^1$H-$^{13}$C NMR. NMR spectra under these experimental conditions showed that ~40 % of the free amino acid was modified (data not shown). The sample was microinjected into one-cell stage zebrafish embryos immediately thereafter.

<u>*Cell lysates from E. coli and SK-N-SH cells.*</u> *E. coli* BL21 (DE3) cells were grown in 200 mL LB medium at 37° C until an $OD_{600}$ of 1.2 was reached. Cells were centrifuged at 5000 r.p.m. for 20 min at 4° C and the supernatant discarded. The pellet was washed with NMR buffer and resuspended in 3 mL of NMR buffer supplemented with 10 % glycerol, 1 mM DTT and 1 % PMSF. We lysed the cells by sonication (Branson, USA) in an ice bath by applying 5 pulses of 10 seconds at 20 W using a sonicator microtip. Between each sonication cycle we waited 2 min to avoid sample heating. Soluble lysate fractions were obtained by centrifugation at 13.000 rpm for 30 min at 4° C. Lysates were aliquoted and stored at -80° C until use.

Human SK-N-SH cells lysates were prepared as previously described.[2] Briefly, 40 million SK-N-SH cells (Sigma, cat.# 86012802) were grown in T175 flasks containing DMEM-Ham's F-12 (PAA laboratories) media supplemented with 10 % FBS (PAA laboratories) at 37° C and 5 % $CO_2$ until they reached 80 % confluence. Cells were detached from the flasks by mild trypsin treatment, washed twice with NMR buffer and resuspended in 1.5 mL NMR buffer supplemented with 1X EDTA-Free complete protease inhibitor cocktail (Roche). Cell lysis was done by 5 freeze/thaw cycles using an ethanol/dry ice bath followed by water bath sonication for 15 min. Lysates were cleared by centrifugation at 13.000 rpm for 30 min at 4° C and the supernatants were aliquoted and stored at -80° C until use. For both *E. coli* and SK-N-SH cell lysates, total protein concentrations were determined using a Bradford assay (BioRad, USA). Repeated thawing and freezing of lysate aliquots were avoided to preserve enzyme activities.[7]

*Reconstituted reduction reactions with pure enzymes and cell extracts.* Reduction reactions with recombinant scMSRA and scMSRB were carried out in triplicates at 10° C for CarMetOx and γ-SynOx and at 25° C for MetOx. In all cases substrate concentration was 100 μM for a reaction volume of 500 μL. Substrate samples were dissolved in NMR buffer at pH 7.0 to which we added 10 mM DTT as electron source[3] and scMSRA (0.4-1 μM) or scMSRB (1.8-3.6 μM). For reactions with endogenous MSRs present in *E. coli* or SK-N-SH cells we adjusted the respective lysate concentrations to 4.0 mg/mL with NMR buffer supplemented with 10 mM DTT and added 100 μM of CarMetOx, γ-SynOx or MetOx. Reactions were carried out in duplicates at 10° C for CarMetOx and γ-SynOx and at 25° C for MetOx. We lowered the reaction temperatures when performing $^{1}H$-$^{15}N$ NMR spectra to minimize amide proton exchange and increase signal-to-noise ratios.

*Zebrafish maintenance.* Handling of zebrafish was done according to national and international guidelines, and carefully monitored by local animal protection authorities. Protocols were approved by the Committee for Ethics of Animal Experiments of the School of Biochemical and Pharmaceutical Sciences of the National University of Rosario. Adult zebrafish were kept at 28.5° C on a 14:10 h light:dark cycle according to standard laboratory procedures.[8] Embryos were staged by morphological features and/or hours post-fertilization (hpf) at 28.5° C according to Kimmel and co-workers.[9] AB/TU (https://zfin.org) wild-type lines were used.

*Zebrafish lysate preparation.* We collected zebrafish embryos at 512-cell and 26-somite stages, washed them three times with NMR buffer and stored them at -80° C until further processing. Frozen embryos (500 for each NMR experiment, twice the number of embryos used for *in vivo* experiments) were homogenized in 1/10 volumes of ice-cold 10X NMR buffer supplemented with EDTA-Free complete protease inhibitor cocktail (Roche) using a Potter-Elvehjem (Thomas, Philadelphia, PA, USA). Homogenates were sonicated by applying 3 pulses of 5 seconds at 10 W using a sonicator microtip (Branson USA). Between each sonication cycle we waited 2 min to avoid sample heating. Extracts were used immediately after preparation. Total protein was 1-2 mg/mL. All steps were performed on ice.

*Zebrafish microinjection.* For NMR experiments in zebrafish embryos, we microinjected the yolk of one-cell stage specimens with 3 nL of 25 mM $^{15}N$-$^{13}C$ isotopically enriched CarMetOx dissolved in 0.3X Danieau's solution[10] using a gas-driven microinjection apparatus (MPPI-2 Pressure Injector, Applied scientific Instrumentation; Eugene, OR, USA). Following established procedures, single rounds of injections were performed within 3 s per embryo.[11] Following injections, embryos were kept in 1X E3 medium in a Petri dish at 28.5° C.[12] In-cell concentrations of microinjected CarMetOx were estimated to be ~ 420 μM, considering that zebrafish embryos are spherical with a diameter of 0.7 mm (~0.18 μL volume).[9] At the desired developmental stage, we collected ~200 embryos with a plastic Pasteur pipette and transferred them into a 5 mm Shigemi tube (Shigemi, Japan) filled with 1X E3 medium and without applying the piston. After 5 min all embryos were settled at the bottom of the tube. We removed excess 1X E3 medium to yield a final volume of ~700 μL, added 70 μL $D_2O$ and recorded NMR experiments. Following data acquisition, embryos were transferred back to a Petri dish with fresh 1X E3 medium, incubated at 28.5° C to evaluate their development and survival rates. Non-injected and CarMetOx-injected embryos were kept as controls to compare with those that were subjected to NMR measurements.

For distribution/localization experiments, 3 nL of a 0.9 mM L-Met-FITC was dissolved in 0.3X Danieau's solution and injected into one-cell stage embryos as described before. The concentration of microinjected L-Met-FITC was ~15 μM. After injection, embryos were incubated in a Petri dish with 1X E3 medium at 28.5° C until they were imaged.

*Zebrafish viability assay.* Embryo survival was evaluated at 26-somite stage on at least three independent samples. As shown in **Figure 4f**, 82.5±4.3 % of CarMetOx injected embryos were alive, whereas 17.5±4.3 % did not survive the injection procedure. Live embryos injected with CarMetOx and subjected to NMR measurements displayed a survival rate of 66.5±9.00 %, whereas 33.5±9.1 % were lost. Survival rates are expressed as mean +/- s.d. It should be noted that in both cases, the reported survival rates included those specimens that were lost due to bad egg quality (~10 % on average) and the general microinjection procedure (~5 % on average), irrespective of the nature of the delivered compound, or whether mock injections were performed. These values were well within the range of general zebrafish microinjection procedures[13]. Delivery of CarMetOx and intracellular exposure reduced viability by a mere ~2.5 %, which strongly suggests that the compound caused no adverse cytotoxicity effects. Collection in NMR tubes and NMR measurements reduced viability by ~16 %, indicating a more pronounced contribution to the overall loss of viability. Therefore, only ~18.5 % of embryos that were lost during the procedure accounted for the combined effects of CarMetOx exposure and general NMR settings. Viability was assessed by bright field microscopy with a Nikon SMZ-1 microscope. We classified embryos as dead when they were necrotic.[14] Embryos inside the NMR tube arrested in development, resembling the previously reported suspended animation state induced by hypoxia.[15] However, normal development resumed after NMR experiments, when embryos were transferred to fresh 1X E3 medium at 28.5° C. Importantly, surviving microinjected embryos continued to develop normally and less than 4 % presented morphological abnormalities, independent of whether they were subjected to NMR experiments, or not. Embryos were imaged again at 48 hpf (long-pec stage). By that stage, embryo mortality remained below 5 %, indistinguishable from non-injected control specimens.

*Bright-field and fluorescence microscopy.* CarMetOx or L-Met-FITC injected-embryos and corresponding controls were observed and imaged at different developmental stages with an Olympus MVX10 fluorescent microscope, a MVPLAPO 1X objective and Olympus DP72 microscope digital camera. Image analysis was done with the DP2-BSW software. Magnifications of 1.6x or 3.2x were used for multiple or single embryos, respectively. For distribution/localization experiments, 3 nL of a 0.9 mM L-Met-FITC was dissolved in 0.3X Danieau's solution and injected into one-cell stage embryos as described before. The concentration of microinjected L-Met-FITC was ~15 μM. After injection, embryos were incubated in a Petri dish with 1X E3 medium at 28.5° C until they were imaged. We used an Olympus U-MGFPHQ/XL mirror unit (DM485HQ dichroic mirror; BP460-480HQ excitation filter; BA495-540HQ barrier filter) on the Olympus MVX10 fluorescence microscope was used. Adobe Photoshop and Illustrator were used for image processing. Embryos were anesthetized with 0.16 mg/ml Tricaine (3-amino benzoic acil ethyl ester; Sigma-Aldrich, A-5040) before imaging.

*NMR spectroscopy.* NMR experiments were recorded on 600 MHz Avance II and 700 MHz Avance III Bruker spectrometers using triple-resonance inverse NMR probes (5mm $^{1}H/D-^{13}C/^{15}N$ TXI) equipped with z-axis, self-shielded gradient coils. 1D and 2D $^{1}H-^{15}N$ NMR spectra for CarMet, CarMetOx or Cametona samples were acquired using SOFAST-HMQC[16] pulse sequences with 60 ms interscan delays. Selective amide proton excitation was achieved with 120° polychromatic PC9 pulses of 3000 μs and 0.01 watts and refocusing using 180° band-selective pulses with uniform responses and phases (REBURP) of 1000 μs and 0.07 watts. Both pulses were centered at 8.5 p.p.m. along the proton dimension. $^{15}N$-decoupling during the acquisition period was accomplished with the Waltz-16 sequence. $^{13}C$ decoupling was accomplished using a smoothed chirp pulse of 500 μs and 45.8 watts. 2D $^{1}H-^{15}N$ SOFAST HMQC spectra were acquired using 256 scans and 1K and 32 complex points for $^{1}H$ and $^{15}N$ dimensions, respectively. Sweep-widths were 16 p.p.m. for $^{1}H$ and 2 p.p.m. for $^{15}N$. As we only detect one signal in CarMet and two in CarMetOx, at similar $^{15}N$ resonance frequencies, we used short sweep-widths and few indirect points without compromising on resolution. This allowed us to save acquisition time and to record more scans for better signal-to-noise ratios within the same experimental time. For time-course experiments, individual NMR spectra were acquired within 35 min. Processing was achieved by zero-filling to 2K and 1K points in $^{1}$H and $^{15}$N, respectively, followed by sine-modulated window function multiplication in both dimensions prior to Fourier transformation and baseline correction.

For $^{1}$H-$^{15}$N SOFAST-HMQC NMR experiments of $^{15}$N enriched γ-Syn and γ-SynOx, we used an interscan delay of 40 ms. Selective amide proton excitation was achieved with 120° polychromatic PC9 pulses of 2600 μs and 0.02 watts and refocusing using 180° band-selective pulses with uniform responses and phases (REBURP) of 1000 μs and 0.12 watts. Both pulses were centered at 8.5 p.p.m. along the proton dimension. $^{15}$N-decoupling during acquisition periods was accomplished with the Waltz-16 sequence. 2D $^{1}$H-$^{15}$N SOFAST HMQC spectra were acquired using 64 scans and 1K and 256 complex points for $^{1}$H and $^{15}$N dimensions, respectively. Sweep-widths were 16 p.p.m. for $^{1}$H and 26 p.p.m. for $^{15}$N. The experimental time for each NMR spectrum during time course experiments was 38 min. Processing was achieved by zerofilling to 2K and 1K points in $^{1}$H and $^{15}$N, respectively, followed by qsine-modulated window function multiplication in $^{1}$H and no apodization in $^{15}$N prior to Fourier transformation and baseline correction.

$^{1}$H-$^{13}$C HSQC spectra of Met and MetOx were acquired using a standard Bruker pulse sequence with sensitivity enhancement (hsqcetgpsisp2.2). We used 16 scans and 2K and 64 complex points for $^{1}$H and $^{13}$C dimensions, respectively. Sweep-widths were 16 pp. for $^{1}$H and 60 p.p.m. for $^{13}$C. Experimental time for each NMR spectrum was 19 min. Processing was done by zero-filling to 2K and 512 points in $^{1}$H and $^{13}$C, respectively, followed by sine-modulated window function multiplication in both dimensions and baseline correction.

1D & 2D $^{1}$H-$^{15}$N SOFAST HMQC spectra of zebrafish embryos injected with CarMetOx were acquired at 15° C. We used a 60 ms interscan delay, 2K scans and 1K and 24 complex points for $^{1}$H and $^{15}$N dimensions, respectively. Sweep-widths were 16 p.p.m. for $^{1}$H and 2 p.p.m. for $^{15}$N. Experimental time for each NMR spectrum was ~3 h. Processing was achieved by zerofilling to 2K and 1k points in $^{1}$H and $^{15}$N, respectively, followed by qsine- and sine- modulated window function multiplication in $^{1}$H and $^{15}$N dimension and baseline correction. After the NMR experiments, we separated the embryos from the surrounding 1X E3 buffer and recorded additional 1D $^{1}$H-$^{15}$N SOFAST HMQC spectra under identical conditions to exclude leakage during the experiment.

We previously calculated that the intracellular concentration of microinjected CarMetOx was ~420 μM. However, this was not directly correlated with the concentration we detected by NMR[17] because spherical embryos did not pack perfectly in the cylindrical NMR tube. About 200 embryos were contained within the receiver-coil volume of the TXI probe (22 mm) occupying a total volume of 36 μL. The inner diameter of $D_2O$ matched Shigemi tubes is 4.2 mm indicating that the effective NMR volume is 304.8 μL, introducing a dilution factor of ~8.5 for microinjected CarMetOx *in vivo* and yielding an effective NMR concentration of ~50 μM. Comparison the 1D $^{1}$H-$^{15}$N SOFAST-HMQC spectra of CarMetOx *in vitro* (conc. 100 μM) and *in vivo* using identical NMR setups showed that we detected most of the microinjected compound, suggesting that it did not bind to intracellular components (figure S4 d).

All spectra were acquired and processed using Topspin 3.5 (Bruker, Biospin). Calibration was performed with DSS.[18] Signal intensities for *in vitro* time-course experiments were obtained for each spectrum using Sparky[19] and plotted vs time in GraphPad Prism to obtain oxidation and reduction curves. In all cases, dead times between sample mixing and the start of acquisition were subtracted.[7] Error bars in the time-course plots represent the average experimental noise of the respective 2D NMR spectra.[20] In-cell reduction profiles were obtained by integrating individual signals in each spectrum and calculating their relative populations with respect to the others. For instance, % *R*-CarMetOx = *R*-

CarMetOx/(*R*-CarMetOx + *S*-CarMetOx + CarMet). We used this strategy to normalize for small experimental variations in the individual experiments.

<u>*Model fitting of MSR activities.*</u> Methionine sulfoxide reduction kinetics were delineated by measuring NMR signal intensities of cross-peaks reporting on the oxidation states of *S*- and *R*- diastereoisomers of Met sulfoxides in γ-SynOx, CarMetOx and the free amino-acid. For γ-SynOx, we selected the Tyr-39 cross-peak and for MetOx the Hγ-Cγ cross-peaks. For γ-SynOx we also considered the sulfoxides of Met-1, which are monitored by the Val-3 cross-peak (*S*- and *R*- overlapped). The concentration of each diastereoisomer was obtained by dividing the initial substrate concentration (100 μM) by 2, which corresponds to the racemic concentrations of *R*- and *S*-sulfoxides obtained upon chemical oxidation with $H_2O_2$ (see **Figure 2** and **S1**). For repair processes of γ-SynOx, we assumed that reduction of Met1 and Met38 sulfoxides pursued independent. Enzymatic parameters were determined by fitting time-course data to the Michaelis-Menten equation using the DynaFit software package.[21] We used the following reaction:

$$\mathrm{MSR} + \mathrm{Met_{OX}} \underset{k_{dMetOX}}{\overset{k_{aMetOX}}{\rightleftharpoons}} \mathrm{MSR\text{-}Met_{OX}} \xrightarrow{k_{cat}} \mathrm{MSR} + \mathrm{Met_{RED}} \quad (1)$$

Where MSR represents scMSRA or scMSRB; $Met_{OX}$ is either CarMetOx, MetOx or γ-SynOx, MSR-$Met_{OX}$ denotes enzyme-substrate complex, $Met_{RED}$ is CarMet, L-Met or γ-Syn, $k_{aMetOX}$ is the association rate constant, $k_{dMetOX}$ the dissociation rate constant and $k_{cat}$ the catalytic rate constant. Differential equations derived from this reaction are:

$$\frac{d[\mathrm{MSR}]}{dt} = -k_{aMetOx}[\mathrm{MSR}][\mathrm{Met_{OX}}] + k_{dMetOx}[\mathrm{MSR\text{-}Met_{OX}}] + k_{cat}[\mathrm{MSR\text{-}Met_{OX}}] \quad (2)$$

$$\frac{d[\mathrm{Met_{OX}}]}{dt} = -k_{aMetOx}[\mathrm{MSR}][\mathrm{Met_{OX}}] + k_{dMetOx}[\mathrm{MSR\text{-}Met_{OX}}] \quad (3)$$

$$\frac{d[\mathrm{MSR\text{-}Met_{OX}}]}{dt} = +k_{aMetOX}[\mathrm{MSR}][\mathrm{Met_{OX}}] - k_{dMetOX}[\mathrm{MSR\text{-}Met_{OX}}] - k_{cat}[\mathrm{MSR\text{-}Met_{OX}}] \quad (4)$$

$$\frac{d[\mathrm{Met_{RED}}]}{dt} = +k_{dMetOx}[\mathrm{MSR\text{-}Met_{OX}}] \quad (5)$$

<u>*Model fitting of Met and γ-Syn oxidation kinetics.*</u> To determine the oxidation rate constants for Met and γ-Syn, we added 100 equivalent (10 mM) of $H_2O_2$ to a 500 μL solution of each substrate (100 μM) dissolved in NMR buffer and started time-course NMR acquisitions. We measured signal intensities of cross-peaks corresponding to the two *S*- and *R*-diastereomers at each time point and fitted curves to the following model:

$$\mathrm{Met_{RED}} + H_2O_2 \xrightarrow{k} \mathrm{Met_{OX}} \quad (6)$$

following a second order reaction where $[H_2O_2]_0 > [Met_{RED}]_0$:

$$\frac{d[\mathrm{Met_{RED}}]}{dt} = -k[\mathrm{Met_{RED}}][H_2O_2] \quad (7)$$

and solving for $[Met_{RED}]$ yield:

$$[\mathrm{Met_{RED}}] = \frac{([H_2O_2]_0 - [\mathrm{Met_{RED}}]_0)\,[\mathrm{Met_{RED}}]_0}{([H_2O_2]_0\, e^{kt([H_2O_2]_0 - [\mathrm{Met_{RED}}]_0)} - [\mathrm{Met_{RED}}]_0)} \quad (8)$$

We fitted experimental data to eq. (8) using non-linear least-squares analysis to obtain the *k* values for *S*- and *R*-diastereoisomers of each substrate. Experiments were performed at 10$^{o}$ C for γ-Syn and at 25$^{o}$ C for Met as previously outlined for reconstituted reduction reactions.

### Supplementary Figures and Tables

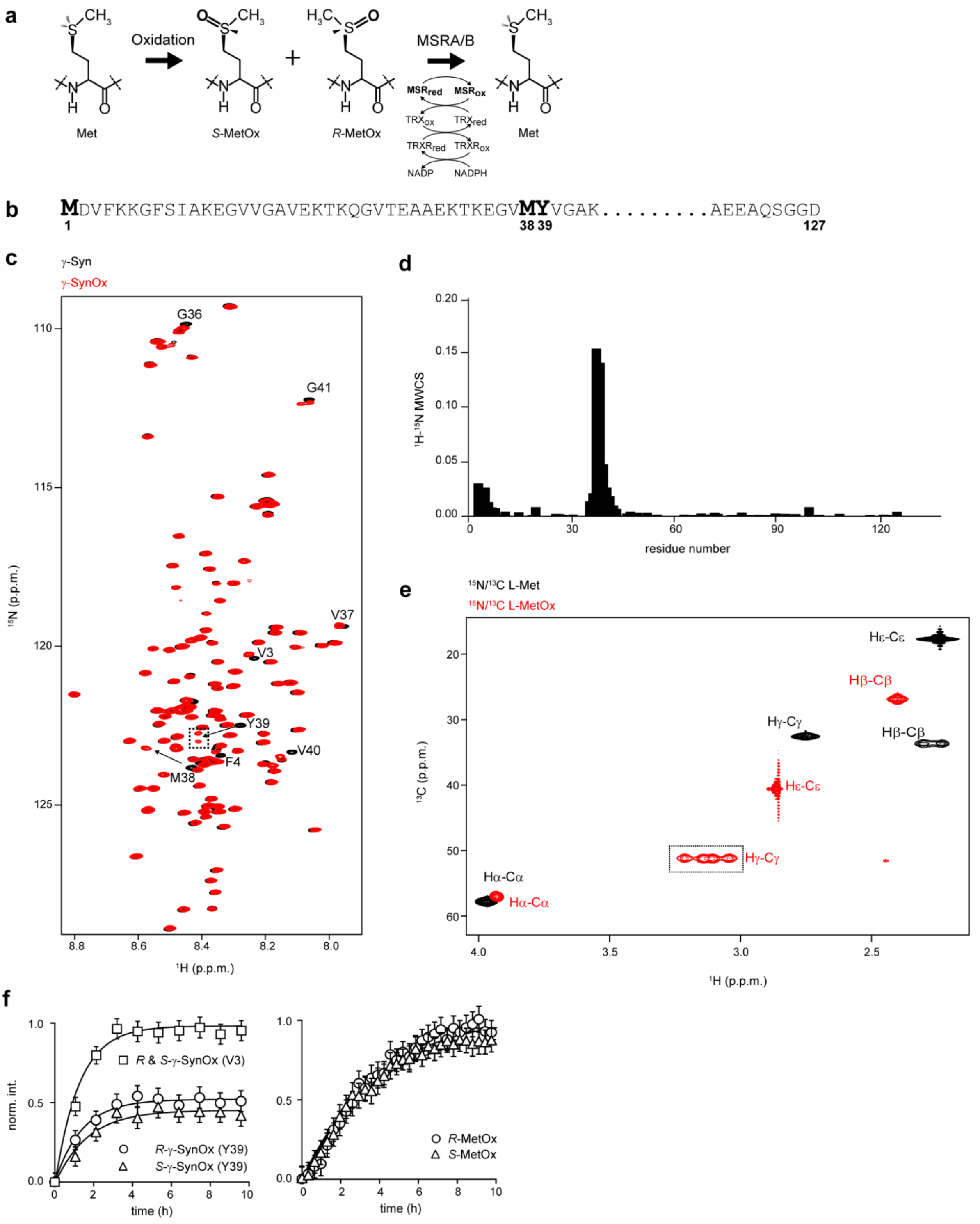


**Figure S1. Oxidation of L-Met and γ-Syn.** (a) Oxidation reaction of Met side-chains to form *S*- and *R*-diastereoisomers of methionine sulfoxide and their stereospecific reduction catalyzed by MSRA and MSRB, respectively. TRX (thioredoxin), TRXR (thioredoxin reductase) and NADPH recycle the active forms of MSRs after each catalytic cycle *in vivo*. (b) Primary sequence of γ-Syn. The position of Met residues and Tyr39 are highlighted. (c) Overlay of 2D $^{1}$H-$^{15}$N SOFAST-HMQC spectra of $^{15}$N isotope-enriched γ-Syn in its reduced (black) and Met-oxidized form (red). Splitting of the Tyr39 cross-peak is induced by *S*- and *R*-diastereomers of Met38, is indicated with a dotted rectangle. (d) γ-Syn $^{1}$H-$^{15}$N mean-weighted chemical shift perturbations in response to Met oxidation. (e) Overlay of $^{1}$H-$^{13}$C HSQC spectra of $^{15}$N-$^{13}$C isotope-enriched reduced (black) and oxidized (red) L-Met. Splitting of the Hγ-Cγ cross-peak corresponding to Met *S*- and *R*-diastereoisomers, is indicated with a dotted rectangle. (f) Time-course oxidation profiles of γ-Syn and Met. Curves show NMR signal intensity buildups of the Tyr39 and Val3 H-N cross-peaks (corresponding to Met1 oxidation) upon addition of $H_2O_2$ (left). For Val3, *R*- and *S*- cross-peaks are overlapped. For the free amino acid, the traces show NMR signal intensity buildups of Hγ-Cγ corresponding to the *R*- and *S*-diastereoisomers of Met plotted as a function of time (right). All experiments were performed with 100 μM γ-Syn or Met in the presence of 10 mM $H_2O_2$. We fitted these data to derive second order stereospecific oxidation kinetics.[22] The values obtained were: (i) γ-Syn Met38, *R*-sulfoxide (*k*= $2.4x10^{-2}$ $M^{-1}$ $s^{-1}$), *S*-sulfoxide (*k*= $1.9x10^{-2}$ $M^{-1}$ $s^{-1}$); (ii) Met1 *R*-and *S*-sulfoxides together (*k*= $2.7x10^{-2}$ $M^{-1}$ $s^{-1}$); (iii) free Met *R*-sulfoxide (*k*= $9.1x10^{-3}$ $M^{-1}$ $s^{-1}$), *S*-sulfoxide (*k*= $9.3x10^{-3}$ $M^{-1}$ $s^{-1}$) in good agreement with previously published oxidation rates. [22]

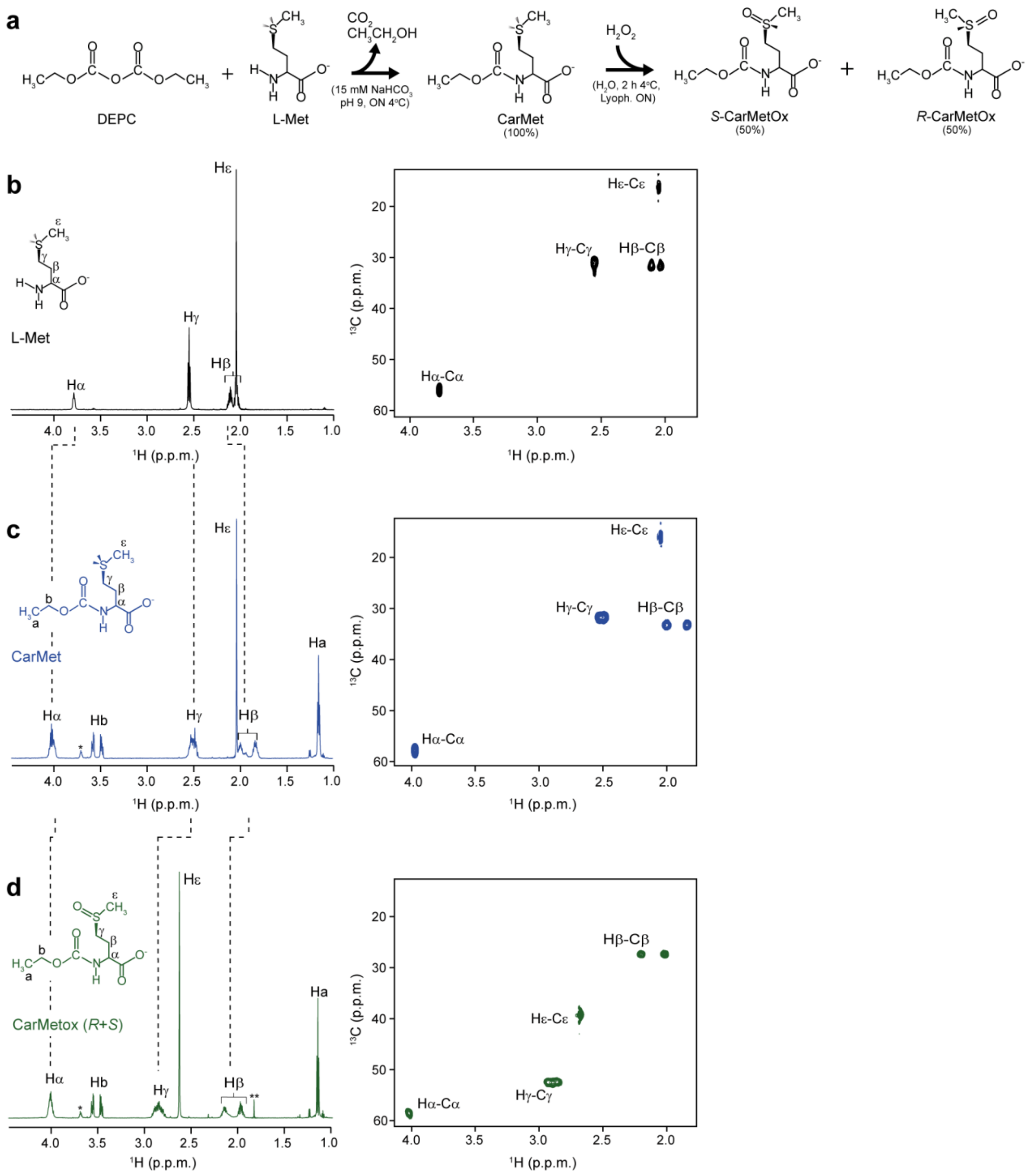


**Figure S2. CarMetOx synthesis.** (a) Reaction steps for synthesizing CarMet and CarMetOx from $^{15}N$-$^{13}C$ isotopically enriched L-Met, non-isotopically enriched DEPC and $H_2O_2$. (b-d) 1D $^1H$ (left) and 2D $^1H$-$^{13}C$ HSQC NMR spectra of L-Met (b), CarMet (c) and CarMetOx (d). 1D $^1H$ NMR spectra were obtained from reaction products using non-isotopically enriched L-Met as starting material. The different atoms present in L-Met and corresponding signals in 1D and 2D NMR spectra are indicated by Greek letters. CarMet and CarMetOx atoms derived from DEPC incorporation are indicated by regular letters. Asterisks in 1D $^1H$ NMR spectra shown in (c) and (d) indicate signals corresponding to impurities of commercial DEPC and $H_2O_2$.

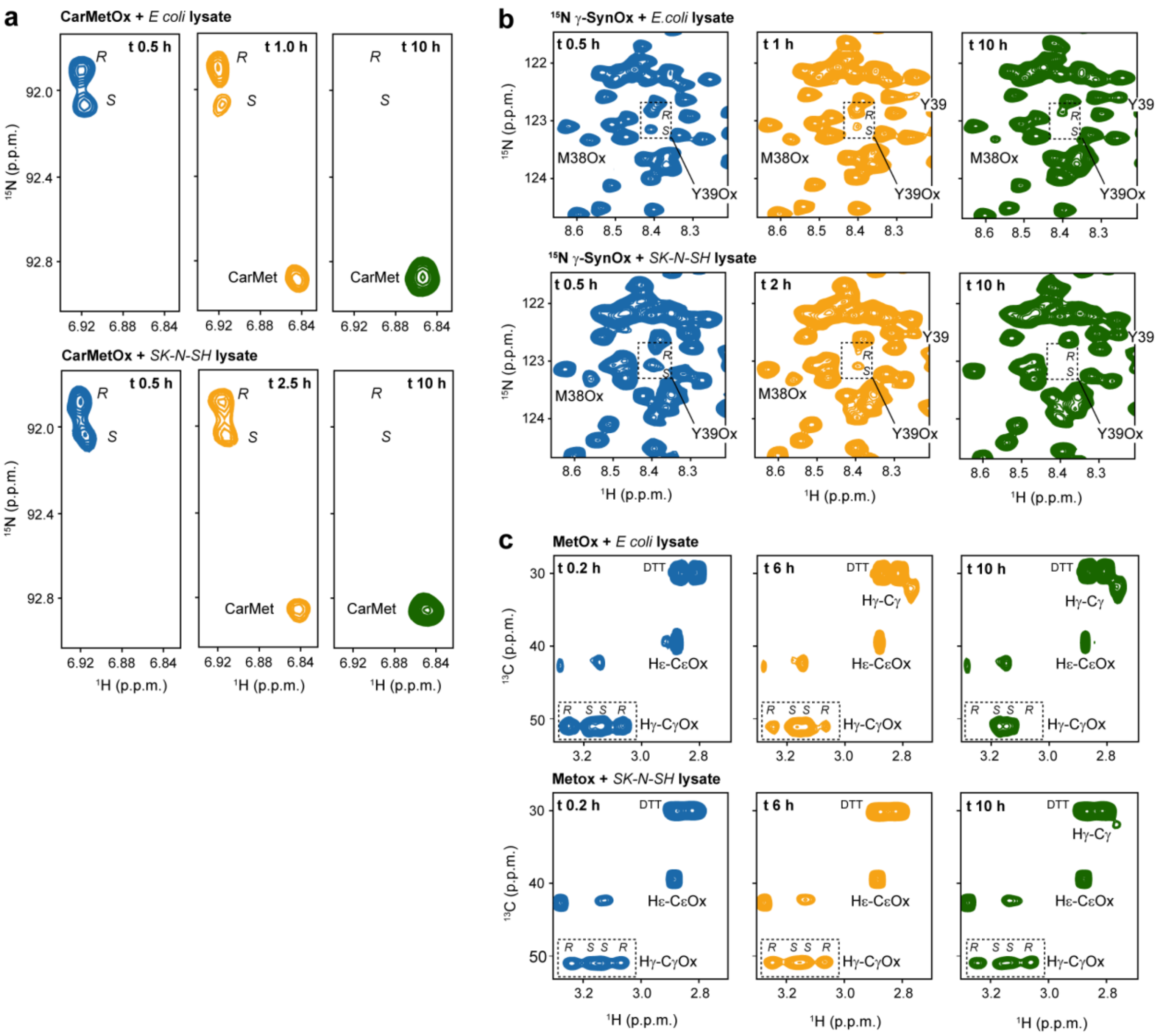


**Figure S3. Stereospecific sulfoxide reductions in *E. coli* and SK-N-SH cell lysates.** NMR monitoring of endogenous MSRs activities on $^{15}$N-$^{13}$C CarMetOx (a), $^{15}$N isotope-enriched γ-SynOx (b), and $^{15}$N-$^{13}$C isotope-enriched MetOx (c), in *E. coli* and SK-N-SH cell lysates. Boxes in (b) and (c) identify the cross-peaks of Tyr39 corresponding to the *S*- and *R*-diastereoisomers of Met38 of γ-SynOx (b) and the $H_\gamma$-$C_\gamma$ cross-peak corresponding to the *S*- and *R*-diastereoisomers of MetOx (c). Experiments were performed by adding 100 μM of CarMetOx, γ-SynOx or MetOx to *E. coli* or SK-N-SH lysates in NMR buffer at pH 7. In all cases, lysates were adjusted to 4 mg/mL of total protein. 10 mM DTT was added as electron source.

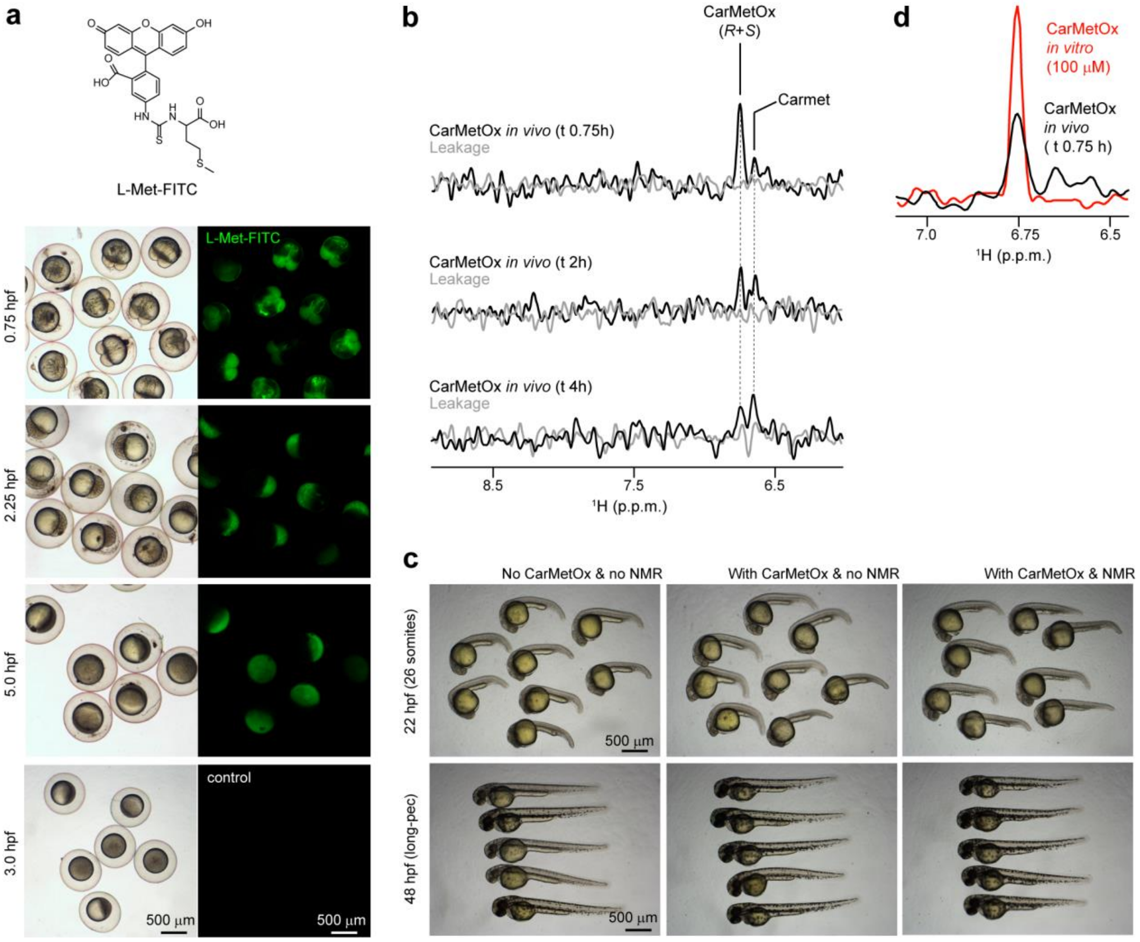


**Figure S4. Zebrafish microinjection controls.** (a) Microscopy of zebrafish embryos microinjected with Met-FITC (top). Bright field (left) and fluorescence (right) microscopy of zebrafish embryos microinjected with FITC-labeled L-Met at different times after delivery. Developmental stages are indicated on the left. (b) 1D $^{1}H$-$^{15}N$ SOFAST-HMQC spectra of CarMetOx injected zebrafish (black) and supernatants (gray) after NMR measurements and removal of embryos. No leakage was detected. Experimental time points are indicated. (c) Bright field microscopy of zebrafish embryos non-injected, CarMetOx-injected or after NMR experiments at the 26-somite (top) and long-pec developmental stages (bottom). (d) 1D $^{1}H$-$^{15}N$ SOFAST-HMQC spectra of CarMetOx injected zebrafish embryos 0.75 h after injection (black) and a 100 μM CarMetOx sample dissolved in NMR buffer at pH 7 (red). Both acquired with identical NMR parameters.

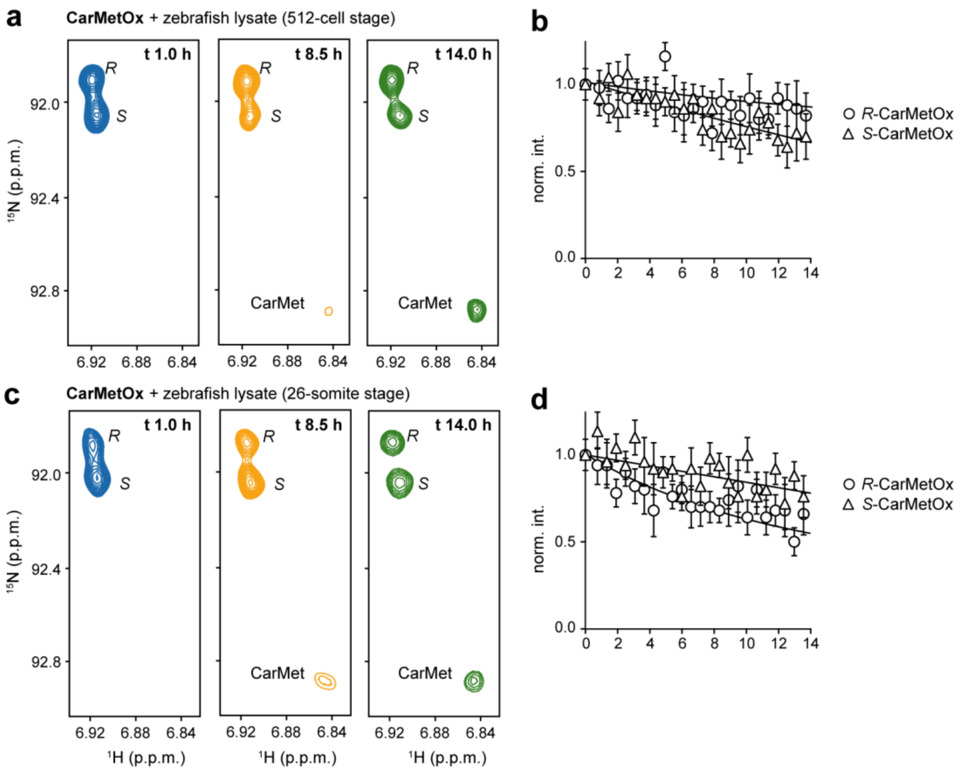


**Figure S5. Stereospecific CarMetOx reduction in zebrafish lysates.** NMR monitoring of endogenous MSRs activities using $^{15}N$-$^{13}C$ CarMetOx in zebrafish lysates prepared from embryos at 512-cell (a, b) and 26-somite stages (c, d). Experiments were performed by adding 100 µM of CarMetOx to lysates in NMR buffer at pH 7. 10 mM DTT was added as electron source.

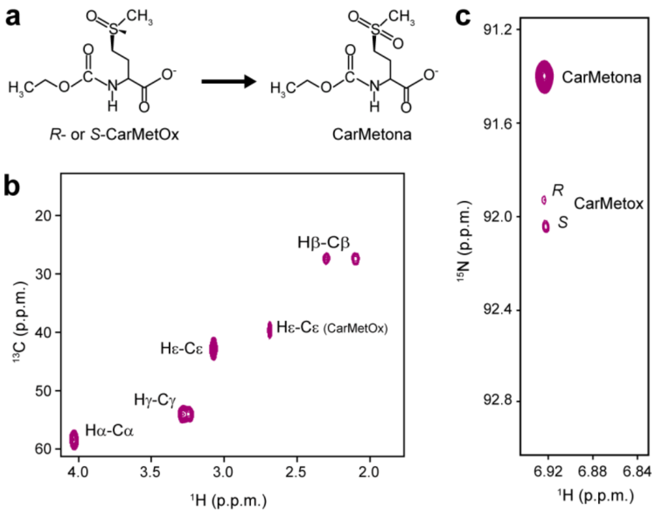


**Figure S6. Spectral features of CarMet sulfone (CarMetona).** (a) Structure of CarMetOx and its further oxidation to CarMetona. $^{1}$H-$^{13}$C HSQC (b) and $^{1}$H-$^{15}$N SOFAST-HMQC (c) spectra of $^{15}$N-$^{13}$C isotopically enriched CarMetona. Cross-peaks corresponding to the different correlations are indicated. A small amount of remnant CarMetOx is indicated in each spectrum. NMR signals of CarMetona do not overlap with those of CarMetOx and are easily identified in $^{1}$H-$^{15}$N correlations, providing a quality control for the absence of sulfones during CarMetOx synthesis.

**Table S1.** Kinetic parameters of CarMetOx, MetOx and γ-SynOx reduction catalyzed by scMSRA and scMSRB.

| Substrate | MSRA | | | MSRB | | |
|---|---|---|---|---|---|---|
| | $k_{cat}$ ($s^{-1}$) | $K_M$ ($\mu M^{-1}$) | $k_{cat}/K_M$ ($s^{-1}$ $M^{-1}$) | $k_{cat}$ ($s^{-1}$) | $K_M$ ($\mu M^{-1}$) | $k_{cat}/K_M$ ($s^{-1}$ $M^{-1}$) |
| **CarMetOx** | 0.5 ± 0.15 | 2 390 ± 720 | **209** | 0.05 ± 0.007 | 1 590 ± 220 | **31** |
| **MetOx** | 1.0 ± 0.2 | 1 631 ± 340 | **613** | 0.052 ± 0.036 | 10 100 ± 7 000 | **5** |
| **γ-SynOx** | 0.97 ± 0.12 | 800 ± 100 | **1 212** | 0.028 ± 0.008 | 980 ± 290 | **28** |

Enzymatic reduction reactions were performed with 100 μM of CarMetOx, γ-SynOx or MetOx in the presence of 0.4-1.0 μM scMSRA and 1.8-3.6 μM scMSRB in NMR buffer at pH 7.0. Experiments with CarMetOx and γ-SynOx were performed at 10° C to maximize NMR signal-to-noise ratios. MetOx reductions were done at 25° C. In all cases, we used 10 mM DTT as electron source.[3]
